# Relaxed 3D genome conformation facilitates the pluripotent to totipotent state transition in embryonic stem cells

**DOI:** 10.1101/2020.11.04.369066

**Authors:** Jiali Yu, Yezhang Zhu, Jiahui Gu, Chaoran Xue, Long Zhang, Jiekai Chen, Li Shen

**Author notes:** These authors contributed equally to this work (Jiali Yu, Yezhang Zhu).

## Abstract

The 3D genome organization is crucial for gene regulation. Although recent studies have revealed a uniquely relaxed genome conformation in totipotent early blastmeres of both fertilized and cloned embryos, how weakened higher-order chromatin structure is functionally linked to totipotency acquisition remains elusive. Using low-input Hi-C, ATAC-seq, and ChIP-seq, we systematically examined the dynamics of 3D genome and epigenome during pluripotency-to-totipotency transition in mouse embryonic stem cells (ESCs). The totipotent 2-cell-embro-like cells (2CLCs) exhibit more relaxed chromatin architecture compared to ESCs, including global weakening of both enhancer-promoter interactions and TAD insulation. While the former leads to inactivation of ESC enhancers and down-regulation of pluripotent genes, the latter may facilitate contacts between the new enhancers arising in 2CLCs and neighboring 2C genes. Importantly, disruption of chromatin loops by depleting CTCF or cohesin promotes ESC to 2CLC transition. Our results thus establish a critical role of 3D genome organization in totipotency acquisition.

**HIGHLIGHTS:**

- Global weakening of the 3D genome conformation during ESC to 2CLC transition
- Loss of enhancer-promoter loops and down-regulation of pluripotent genes in 2CLCs
- Inactivation of ESC enhancers and formation of new enhancers in 2CLCs
- Disruption of chromatin loops by depleting CTCF or cohesin promotes 2CLC emergence

## INTRODUCTION

In mammals, intense chromatin remodeling occurs shortly after fertilization, which involves both reprogramming of epigenetic modifications and reorganization of chromatin architecture (Xu and Xie, 2018; Xu et al., 2020). This process prepares both maternal and paternal genomes for the zygotic genome activation (ZGA), which takes place at the late 1-cell and 2-cell (2C) stages in mouse embryos and is characterized by transient expression of a group of 2C-specific genes and repeats, such as *Zscan4* cluster genes and murine endogenous retrovirus with leucine tRNA (MERVL) repeats (Falco et al., 2007; Hamatani et al., 2004; Peaston et al., 2004). It is noteworthy that ZGA is coupled with the acquisition of totipotency, the ability of a cell to generate all cell types in an organism, including both the embryo proper and extraembryonic tissues. In contrast, embryonic stem cells (ESCs), which are derived from the inner cell mass (ICM) of the blastocyst-stage embryos, are considered pluripotent, as they can differentiate into all three germ layers and germ cells of the embryo but rarely into extraembryonic tissues. Intriguingly, a rare dynamic subset of cells in mouse ESC cultures has been discovered to possess the ability to contribute to both embryonic and extraembryonic tissues as well, showing a totipotent-like developmental potency similar to the 2C-stage blastomeres and highly expressing 2C-specific transcripts (Macfarlan et al., 2012). These cells are thus known as 2C-like cells (2CLCs) and have been demonstrated to be a unique model for understanding totipotency and ZGA. Nevertheless, it remains elusive what defines the chromatin state of a totipotent cell.

Proper 3D genome architecture is crucial for gene regulation. It has been reported that the higher-order chromatin structures of mouse zygotes and 2C embryos are remarkably relaxed, with weakened topologically associating domains (TADs) compared to those of later-stage embryos and somatic cells (Du et al., 2017; Ke et al., 2017). Similar relaxed chromatin architectures of early embryos were also noted in other organisms including fly, fish, and human (Chen et al., 2019; Hug et al., 2017; Kaaij et al., 2018). Moreover, when a somatic nucleus is reprogramed to the totipotent state during somatic cell nuclear transfer (SCNT), its chromatin architecture becomes markedly relaxed as well (Zhang et al., 2020). It appears that the relaxed genome architecture is a unique feature of totipotency. Indeed, the totipotent 2CLCs have also been reported to exhibit increased histone mobility and chromatin accessibility as compared to ESCs (Eckersley-Maslin et al., 2016; Ishiuchi et al., 2015). However, the higher-order chromatin structure in 2CLCs has not been studied genome wide, and how relaxed genome architecture is functionally linked to totipotency remains elusive. In this study, we profiled the dynamics of both 3D genome and epigenome during the spontaneous pluripotent to totipotent state transition in mouse ESCs and defined the role of chromatin architecture relaxation in establishing totipotency.

## RESULTS

### Global weakening of the 3D genome conformation during ESC to 2CLC transition

To investigate the 3D genome architecture in 2CLCs, we generated a mouse ESC line containing a tdTomato transgene driven by the MERVL promoter (MERVL::tdTomato), which has been established as a faithful indicator of the 2C-like state (Macfarlan et al., 2012). Consistent with previous reports, about 0.4% of tdTomato+ cells (i.e., 2CLCs) were observed in this ESC line by flow cytometry (Figure S1A). We then purified tdTomato+ 2CLCs and tdTomato- ESCs and performed RT-qPCR and RNA-seq analyses to validate the fidelity of the purified 2CLCs (Figures S1B-S1H). We found in 2CLCs 1,105 up-regulated genes including reported 2C-specific genes and 244 down-regulated genes (Figure S1B). After validating the purified cells, we carried out small-scale *in situ* Hi-C (sisHi-C) (Du et al., 2017) to determine the chromatin conformation landscapes in 2CLCs and ESCs. About 200 million valid read pairs were obtained for both 2CLCs and ESCs, and the biological replicates were highly correlated (Table S1).

We first examined large-scale chromosome folding by examining active (A) and inactive (B) compartmentalization of the genome. As shown by contact maps and principal component 1 (PC1) values (Lieberman-Aiden et al., 2009), A/B compartments are largely maintained during ESC to 2CLC transition (Figures 1A and 1B). Nevertheless, we observed significant decrease in the strength of compartmentalization in 2CLCs compared to ESCs (Figure 1C). Contacts between B compartments decreased and interaction frequency between A/B compartments increased (Figures S2A and S2B). In addition, we identified 7.9% of genomic intervals with A-to-B (1.0%) or B-to-A (6.9%) compartment switching during ESC to 2CLC transition (Figure S2C). As expected, total transcription within A-to-B switching compartments was down-regulated, while B-to-A switching compartments showed increased transcription during ESC to 2CLC transition (Figure S2D). Although we found a few 2C-specific genes such as the *Tdpoz* family genes in the B-to-A switching compartments (Figure S2E), it is worth noting that only a small fraction (92/1,105, 8.3%) of 2C-specific genes manifested B-to-A switching (Figure S2F), indicating that compartment switching is not a general requirement for the activation of 2C-specific genes in 2CLCs. Interestingly, genomic intervals showing A-to-B switching are slightly enriched for previously identified ESC enhancers (Hnisz et al., 2013) (Figure S2G), implying inactivation of ESC enhancers in 2CLCs.

**Figure 1.**
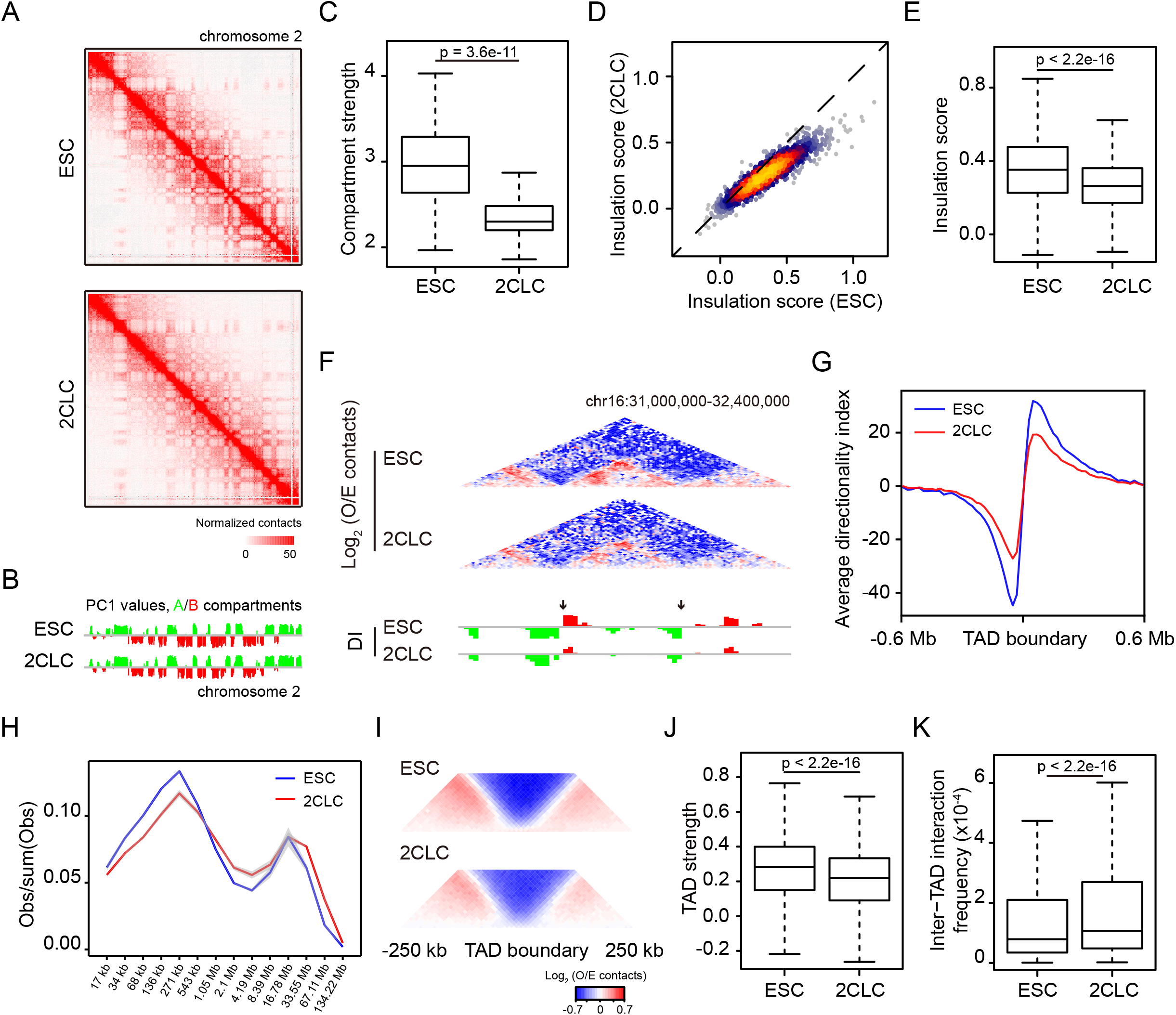
Global weakening of the 3D genome conformation during ESC to 2CLC transition. (**A**) Hi-C contact maps at 250-kb resolution across the entire chromosome 2. (**B**) PC1 values at 500-kb resolution across the entire chromosome 2. (**C**) Box plot showing compartment strength in ESCs and 2CLCs. (**D**) Scatter plot comparing insulation scores at ESC TAD boundaries (Dixon et al., 2012) between ESCs and 2CLCs. (**E**) Box plot showing the insulation score at TAD boundaries. Note that higher score denotes higher insulation potential. (**F**) Genome browser shot showing Hi-C contacts (top) and directionality index (DI) (bottom) at 20-kb resolution. Arrows indicate TAD boundaries. O/E, observed/expected. (**G**) Average DI in a 0.6 Mb region centered on TAD boundaries. (**H**) Contact probability as a function of the genomic distance in logarithmic bins. Lines represent means of biological replicates; edges of semi-transparent ribbons represent individual data points of the two biological replicates. (**I**) Aggregate Hi-C contact maps around TAD boundaries. (**J**) Box plot comparing TAD strengths in ESCs and 2CLCs. (**K**) Box plot showing inter-TAD interaction frequencies in ESCs and 2CLCs.

We next asked whether the local insulation of TADs was reprogrammed during ESC to 2CLC transition. Examination of insulation scores (Crane et al., 2015) at the previously reported ESC TAD boundaries (Dixon et al., 2012) revealed significant attenuation of TAD insulation in 2CLCs as compared to ESCs (Figures 1D, 1E, and S2H), which is further supported by genome-wide decrease of absolute directionality index (Dixon et al., 2012) in 2CLCs (Figures 1F and 1G). In line with these observations, global contact decay curves showed reduced frequencies of short-to intermediate-range interactions (<500 kb) but increased frequencies of long-range interactions (>1 Mb) in 2CLCs as compared to ESCs (Figure 1H). Indeed, 2CLCs exhibited weaker TAD strength and higher frequencies of inter-TAD interactions (Figures 1I-1K). Despite the global reduction of TAD insulation, the numbers of TADs called in 2CLCs (n= 3,674) and ESCs (n= 3,625) were comparable, of which the vast majority (n= 2,957) overlapped with each other (Figure S2I), supporting the overall conserved TADs among different cell types and even across species (Dixon et al., 2012; Rao et al., 2014). Taken together, our observations suggested a genome-wide weakening of TAD insulation in 2CLCs instead of specific gain or loss of TADs.

### Transcription of 2C-specific genes and repeats did not establish local chromatin insulation in 2CLCs

Active transcription was reported to be frequently correlated with chromatin insulation (Bonev et al., 2017; You et al., 2020). Given that 2CLCs highly express a set of 2C-specific genes and the MERVL repeats, we asked whether active transcription at these loci could establish novel insulation in 2CLCs. However, we did not observe any increase in insulation scores at the promoters of up-regulated genes in 2CLCs, indicating no significant changes in chromatin insulation around 2C genes (Figure S2J). Similarly, the highly expressed MERVL repeats also did not cause increased local chromatin insulation in 2CLCs (Figure S2K). Thus, our results support previous observations that transcription is not sufficient to cause chromatin insulation (Bonev et al., 2017).

### Loss of enhancer-promoter interactions and down-regulation of pluripotent genes during ESC to 2CLC transition

Reduced TAD insulation indicated loss of chromatin loops. Indeed, we observed in 2CLCs a mild loss of total chromatin loops previously identified in ESCs, most of which are anchored at TAD boundaries (Rao et al., 2014) (Figures 2A and 2B). Given that chromatin loops not only link TAD boundaries but also tether enhancers and promoters, we specifically analyzed interactions between reported ESC enhancer-promoter pairs (Kieffer-Kwon et al., 2013). This analysis revealed that enhancer-promoter loops were also reduced in 2CLCs (Figures 2C and 2D). Furthermore, reanalysis of published Hi-C datasets showed reduced strength of both types of chromatin loops in 2C embryos as compared to ICM (Figures S2L and S2M). These observations suggested that loss of chromatin loops, both between TAD boundaries and between enhancer-promoter pairs, is a unique feature of totipotency. Consistent with loss of enhancer-promoter interactions, we found that down-regulated genes in 2CLCs were located closer to ESC enhancers (Figure 2E) and were enriched for ESC super-enhancer neighboring genes (Figure 2F). These super-enhancer neighboring genes also displayed significantly lower expression in 2CLCs (Figure 2G). In line with the critical role of ESC super-enhancers in regulating pluripotent genes (Whyte et al., 2013), down-regulated genes in 2CLCs were functionally enriched for pluripotency (Figure S1H). Indeed, almost all pluripotent genes we examined were down-regulated in 2CLCs, including *Pou5f1*, *Sox2*, *Nanog*, *Myc*, *Klf4*, *Esrrb*, *Lin28a*, and *Rex1* (Figures 2H and 2I), likely due to weakened enhancer-promoter interactions in 2CLCs (Figures S3A and S3B).

**Figure 2.**
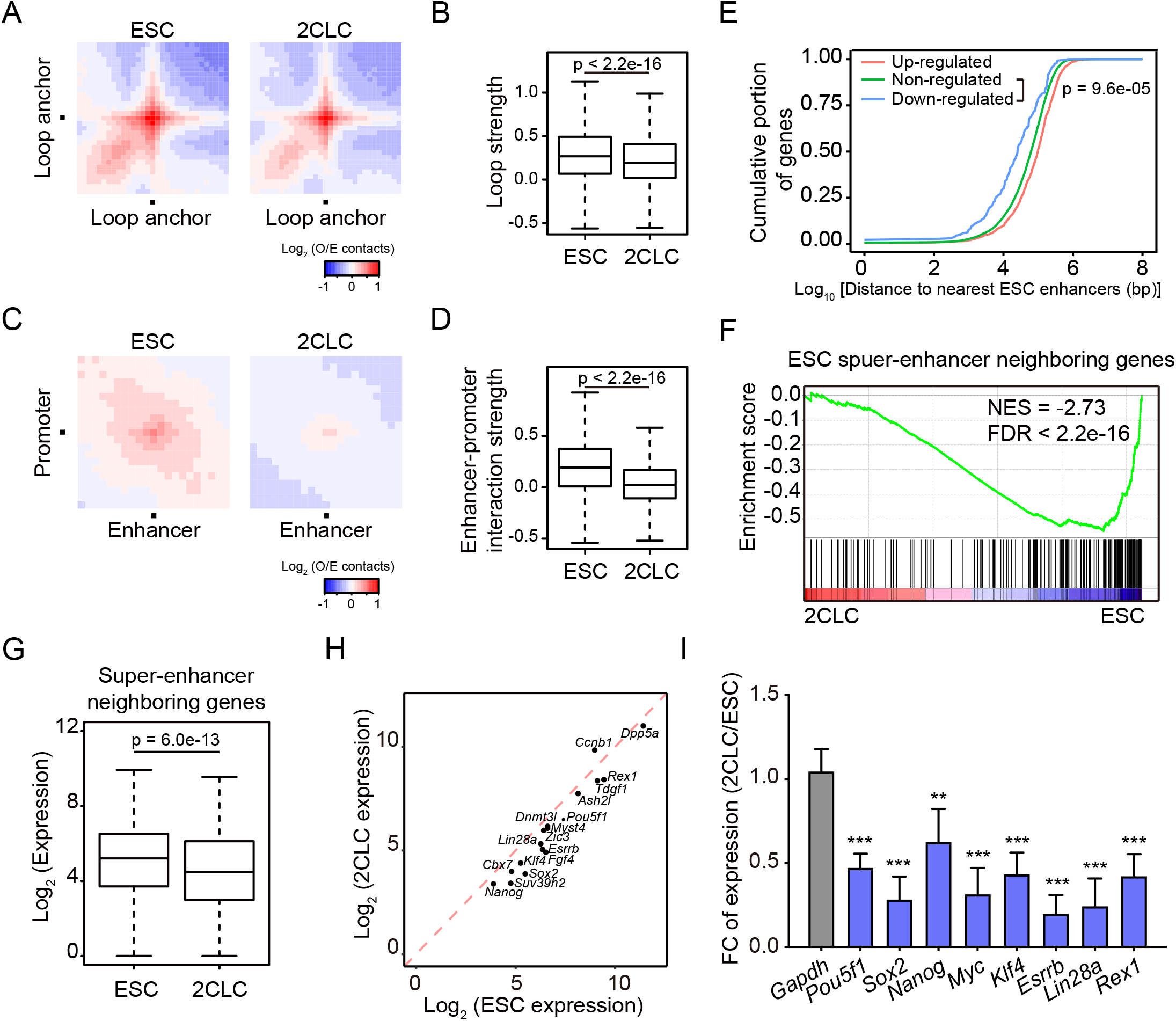
Loss of enhancer-promoter interactions and down-regulation of pluripotent genes in 2CLCs. (**A**) Aggregate Hi-C contact maps between pairs of loop anchors (Rao et al., 2014). (**B**) Box plot showing loop strengths in ESCs and 2CLCs. (**C**) Aggregate Hi-C contact maps between ESC enhancer-promoter pairs (Kieffer-Kwon et al., 2013). (**D**) Box plot showing enhancer-promoter interaction strengths in ESCs and 2CLCs. (**E**) Down-regulated genes in 2CLCs tend to locate closer to ESC enhancers. Shown are cumulative frequency curves comparing the distances between transcription start sites (TSS) of non-/up-/down-regulated genes and their nearest ESC enhancers. P value was calculated using two-sided Kolmogorov–Smirnov test. (**F**) Gene set enrichment analysis indicating that down-regulated genes in 2CLCs were enriched for ESC super-enhancer neighboring genes. Red, up-regulated genes in 2CLCs; blue, down-regulated genes in 2CLCs. (**G**) Box plot comparing the expression of super-enhancer neighboring genes in ESCs and 2CLCs. (**H**) Scatter plot comparing expression of pluripotent genes in ESCs and 2CLCs. (**I**) Relative expression levels of pluripotent genes in 2CLCs versus ESCs by RT-qPCR. Data are normalized to *Actin* and are presented as mean ± SD, **p < 0.01, ***p < 0.001 (Multiple *t* tests). *Gapdh* serves as a control.

### Inactivation of ESC enhancers and formation of new enhancers in 2CLCs

Chromatin accessibility is a strong indicator of the regulatory activity of enhancers (Thurman et al., 2012). To examine whether loss of enhancers-promoters loops in 2CLCs affected enhancer activities, we carried out ATAC-seq analysis (Buenrostro et al., 2013) for both ESCs and 2CLCs. As expected, chromatin accessibility around MERVL repeats were strongly induced in 2CLCs (Figure 3A), consistent with the activation of MERVL during ESC to 2CLC transition. In contrast, meta-analysis at ESC enhancers and super-enhancers revealed a drastic decrease of chromatin accessibility in 2CLCs (Figure 3B), indicating inactivation of ESC enhancers, particularly super-enhancers, during the pluripotent to totipotent state transition. In agreement with this observation, promoters of ESC super-enhancer neighboring genes and down-regulated genes also displayed reduced chromatin accessibility (Figure 3C). To further confirm that ESC enhancers were silenced during ESC to 2CLC transition, we performed ultra-low-input native ChIP-seq (ULI-NChIP-seq) (Brind’Amour et al., 2015) and compared genome-wide histone modifications between ESCs and 2CLCs, including H3K27Ac, H3K4me1, H3K4me3, H3K27me3, and H3K9me3 (Figures 3A-3C, S3C, S3D, and Table S1). As expected, MERVL repeats exhibited remarkable increase in H3K27Ac, H3K4me1, and H3K4me3 in 2CLCs (Figure 3A). However, ESC enhancers and super-enhancers displayed dramatic decrease in both H3K27Ac and H3K4me1 (Figure 3B), confirming the inactivation of ESC enhancers during ESC to 2CLC transition. Consistently, ESC enhancers are also inactive in 2C embryos and are established gradually during preimplantation development (Figure S4A).

**Figure 3.**
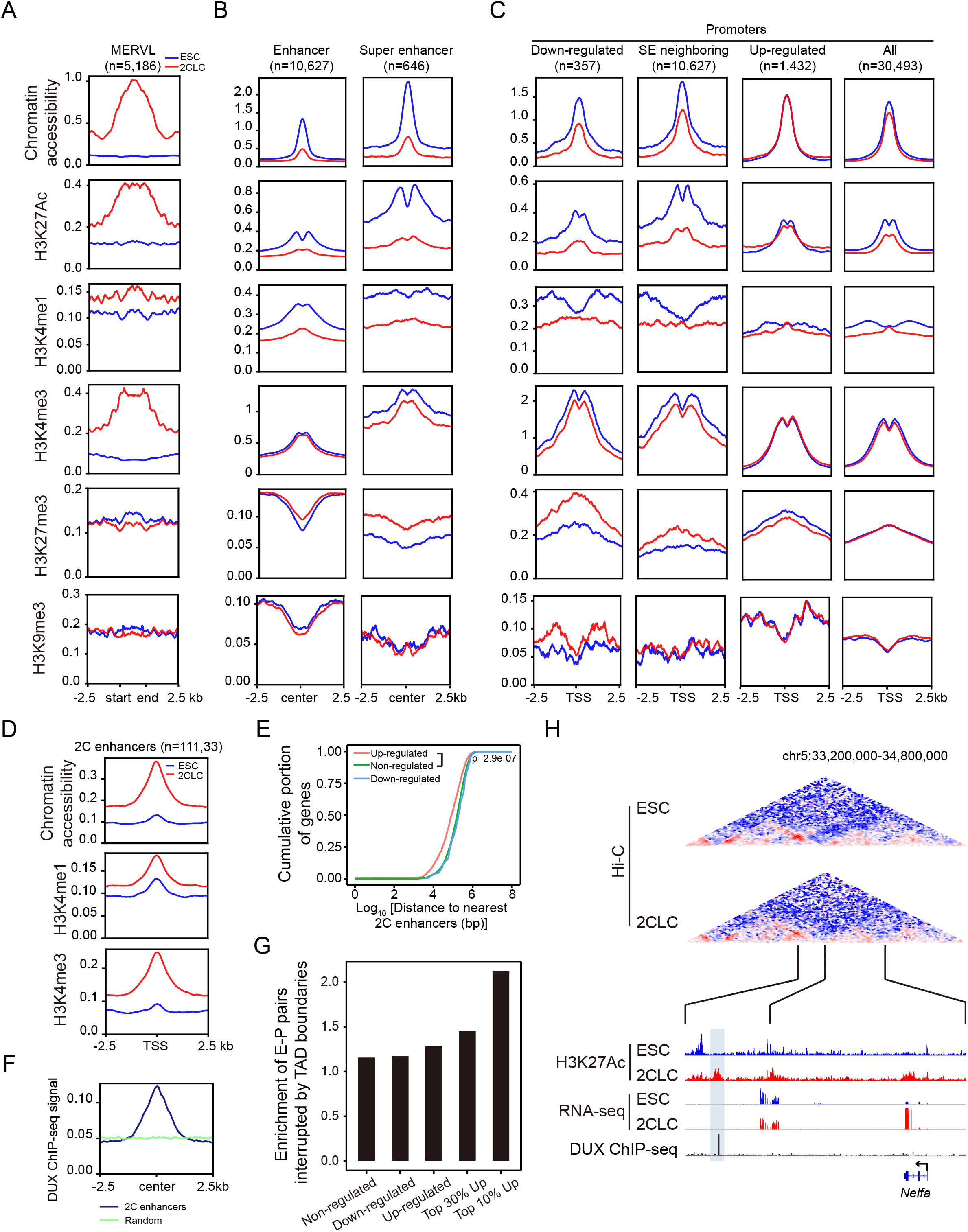
Inactivation of ESC enhancers and formation of new enhancers in 2CLCs. (**A-C**) Average chromatin accessibility, H3K27Ac, H3K4me1, H3K4me3, H3K27me3 and H3K9me3 signals around MERVL (A), ESC enhancers/super-enhancers (B), and indicated promoters (C) in ESCs and 2CLCs. SE, super-enhancer. (**D**) Average chromatin accessibility, H3K4me1 and H3K4me3 signals around 2C enhancers. (**E**) Up-regulated genes in 2CLCs tend to locate closer to 2C enhancers. Shown are cumulative frequency curves comparing the distances between TSS of non-/up-/down-regulated genes and their nearest 2C enhancers. P value was generated using two-sided Kolmogorov–Smirnov test. (**F**) Average DUX ChIP-seq signals around 2C enhancers. Equal number of random regions were used as the control. (**G**) Bar plot showing that up-regulated genes tend to have ESC TAD boundaries separating them from neighboring 2C enhancers. Values represent enrichment of boundary presence between the promoters of indicated genes and their nearest 2C enhancers. Boundary presence between the same promoters and the symmetrical sites of their nearest 2C enhancers serves as the control. (**H**) Genome browser shot showing a TAD boundary between the promoter of *Nelfa* and a 2C enhancer (highlighted). Shown are Hi-C contact maps at 20-kb resolution (top) and genome browser tracks of H3K27Ac ChIP-seq, RNA-seq and DUX ChIP-seq signals (bottom).

Among the five histone modifications we profiled, H3K27Ac peaks displayed the most notable difference in genome-wide distribution between ESC and 2CLCs, with more peaks enriched in distal intergenic regions in 2CLCs (Figure S3D). Examination of these 2CLC-specific distal H3K27Ac peaks (n=11,133) not only revealed increases of chromatin accessibility, H3K27Ac, H3K4me1 and H3K4me3 during ESC to 2CLC transition (Figure 3D), but also high levels of chromatin accessibility and H3K27Ac in 2C embryos (Figure S4A), suggesting the formation of new enhancers (i.e., 2C enhancers) at these regions during totipotency acquisition. Indeed, up-regulated genes in 2CLCs were located closer to these 2C enhancers in the genome (Figure 3E). We found strong DUX ChIP-seq signals at these 2C enhancers, suggesting that they may arise from DUX binding (Figure 3F). Interestingly, the most up-regulated genes during the 2C-like transition tend to have ESC TAD boundaries separating them from neighboring 2C enhancers (Figure 3G). Therefore, weakened TAD insulation may facilitate contacts between the 2C enhancers activated by sporadically expressed DUX in ESCs and the promoters of their neighboring 2C genes, as exemplified by *Nelfa*, an early driver of the 2C-like state (Hu et al., 2020) (Figure 3H). Taken together, inactivation of ESC enhancers, formation of 2C enhancers, and weakening of TAD boundaries in 2CLCs function synergistically to fully activate the 2C-like transcriptional program.

### Depletion of CTCF or cohesin complex facilitates the ESC to 2CLC transition

Our results have demonstrated that 2CLCs possess a relaxed chromatin architecture, including weakened TAD boundary loops and reduced enhancer-promoter loops. While reduced enhancer-promoter loops results in down-regulation of pluripotent genes, weakened TAD insulation may facilitate the activation of 2C genes by sporadically activated 2C enhancers (Figure S4B). We thus hypothesized that disruption of chromatin loops by depletion of CTCF or the cohesin complex would facilitate the ESC to 2CLC transition. To directly test this, we knocked down *Ctcf* and core cohesin complex genes including *Smc1a*, *Smc3* and *Rad21* individually with small interfering RNAs (siRNAs) in MERVL::tdTomato ESCs. As a control, we also knocked down *Yy1*, a structural regulator that was reported to mediate only enhancer-promoter interactions but not TAD boundary loops (Weintraub et al., 2017). Supporting our hypothesis, depletion of either CTCF or cohesin, but not *Yy1*, significantly increased the fraction of tdTomato+ cells (Figure 4A), as well as the expression of MERVL and 2C-specific genes including *Dux*, *Zscan4d*, *Zfp352* and *Tdpoz4* (Figure 4B), demonstrating an enhanced ESC to 2CLC transition upon disruption of all chromatin loops rather than enhancer-promoter loops alone. We further examined the effect of CTCF and cohesin removal on all 2C genes by reanalyzing published RNA-seq datasets of ESCs with acute protein degradation of CTCF or RAD21 (Nora et al., 2017; Zhang et al., 2020). This analysis also showed that depletion of either CTCF or cohesin could result in activation of 2C genes (Figures 4C and 4D).

**Figure 4.**
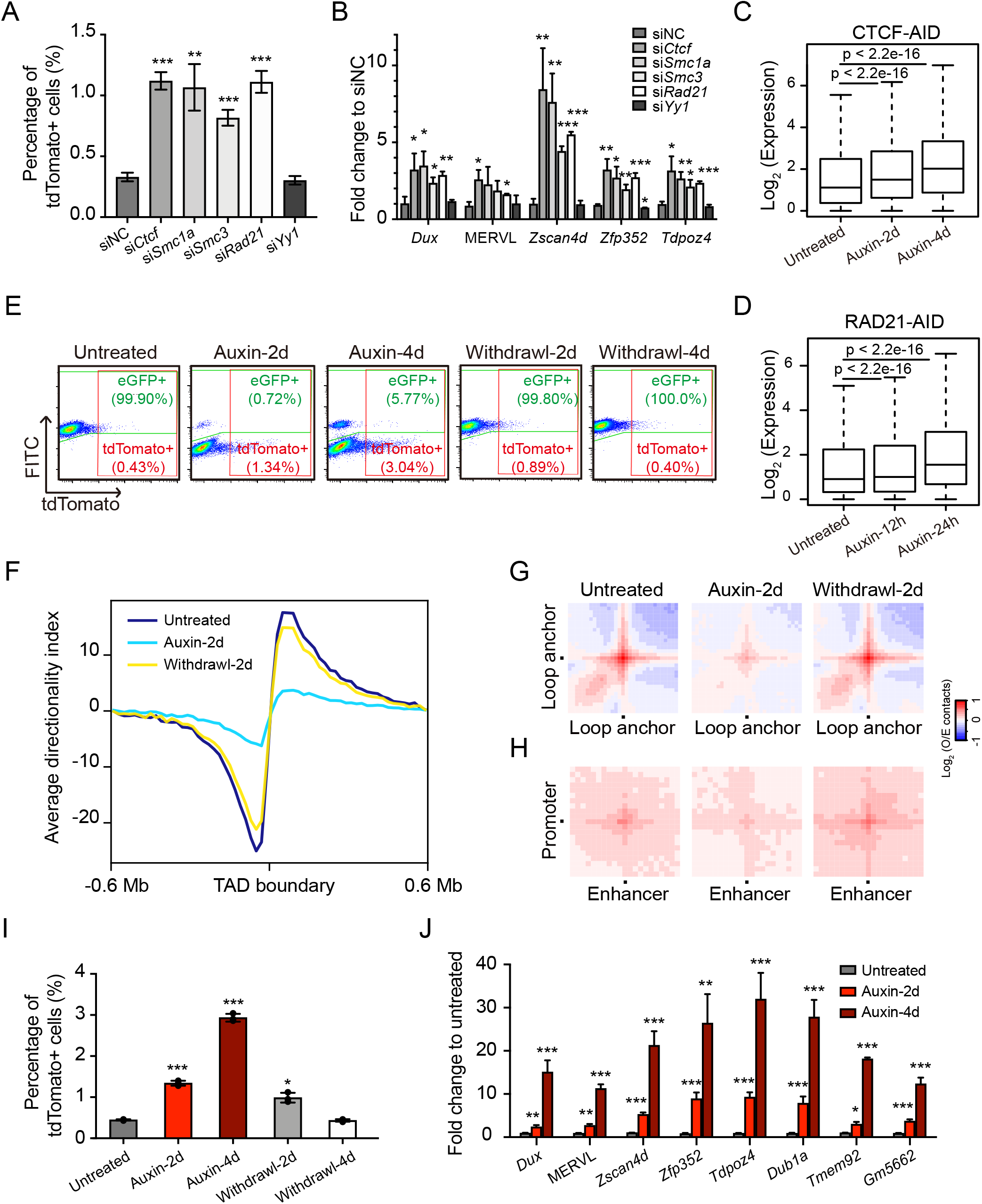
Depletion of CTCF or cohesion complex facilitates ESC to 2CLC transition. (**A**) The percentage of 2CLCs after knocking down *Ctcf*, *Smc1a*, *Smc3*, *Rad21*, or *Yy1*. Data are presented as mean ± SD, n=3. **p < 0.01, ***p < 0.001 (Multiple *t* tests). (**B**) Relative expression levels of 2C-specific transcripts after knocking down *Ctcf*, *Smc1a*, *Smc3*, *Rad21*, or *Yy1*. Data are presented as mean ± SD, n=3. *p < 0.05, **p < 0.01, ***p < 0.001 (Multiple *t* tests). (**C-D**) Box plot showing up-regulation of 2C-specific genes upon acute depletion of CTCF (C) or RAD21 (D) protein by auxin-inducible degron (AID) system in ESCs. Analyses were performed using published RNA-seq datasets (Nora et al., 2017; Zhang et al., 2020). (**E**) Representative FACS analysis of 2C-CTCF-AID cells treated with or without auxin. Note that endogenous CTCF in this cell line is fused with an eGFP tag to indicate CTCF protein levels. (**F**) Average DI in a 0.6 Mb region centered on TAD boundaries. (**G**) Aggregate Hi-C contact maps between pairs of loop anchors. (**H**) Aggregate Hi-C contact maps between ESC enhancer-promoter pairs. (**I**) The percentage of 2CLCs after auxin treatment. Data are presented as mean ± SD, n=3. *p < 0.05, ***p < 0.001 (Multiple *t* tests). (**J**) Relative expression levels of 2C-specific transcripts after auxin-induced CTCF degradation. Data are presented as mean ± SD, n=3. *p < 0.05, **p < 0.01, ***p < 0.001 (Multiple *t* tests).

We further constructed an ESC line (2C-CTCF-AID) containing both the MERVL::tdTomato reporter and an auxin-inducible degron (AID) system targeting CTCF. Endogenous CTCF in this cell line is also fused with an eGFP tag to indicate CTCF protein levels (Nora et al., 2017). As shown by flow cytometry analysis, 2 days of auxin treatment led to a complete depletion of the CTCF protein, which can be fully restored 2 days after auxin withdrawal (Figure 4E). Reanalyzing published Hi-C data (Nora et al., 2017) confirmed weakened TAD insulations together with reduced chromatin loops upon 2 days of auxin treatment (Figures 4F-4H). Similar to *Ctcf* knockdown, the percentage of tdTomato+ 2CLCs in the culture (Figures 4E and 4I) as well as the expression of MERVL and 2C-specific genes (Figure 4J) dramatically increased upon CTCF degradation. It is noteworthy that CTCF-depletion-induced increase in ESC to 2CLC transition is reversible, as the percentage of tdTomato+ 2CLCs returned to the initial state after auxin withdrawal and CTCF recovery (Figures 4E and 4I). Thus, our results strongly suggested that the higher-order chromatin structure in ESCs is an impediment to the 2C-like state transition.

## DISCUSSION

In addition to epigenetic modifications, 3D chromatin architecture has been increasingly recognized as another regulatory layer of gene expression, which is tightly linked to cell identity. ESC differentiation has been shown to coincide with formation or strengthening of TAD boundary loops, indicating a relatively relaxed chromatin architecture in ESCs compared to differentiated cells (Bonev et al., 2017; Pekowska et al., 2018). Furthermore, mouse zygotes and 2C embryos also display more relaxed chromatin architectures compared to later-stage embryos, with much weaker TAD boundaries (Du et al., 2017; Ke et al., 2017). Similar relaxed chromatin architectures of early embryos were also observed in other organisms including fly, fish, and human (Chen et al., 2019; Hug et al., 2017; Kaaij et al., 2018). Therefore, high developmental potency appears to be associated with relaxed higher-order chromatin structure. In this study, using the pluripotency-to-totipotency transition in mouse ESCs as a model, we defined the role of genome architecture relaxation in establishing totipotency. This spontaneous transition represents a simplified reprogramming process in totipotency acquisition, which is valuable to investigate the functional contribution of one single factor among the complex developmental events. But the scarcity of spontaneous 2C-like transition in the ESC culture limits such investigations. By using low-input technologies, we presented genome-wide chromatin contact maps of totipotent 2CLCs and pluripotent ESCs, together with their transcriptome, chromatin accessibility, and histone modification profiles. This rich dataset has allowed us to investigate the dynamics of chromatin state and gene expression during the totipotent to pluripotent state transition.

We revealed that chromatin architecture of ESCs became more relaxed during the spontaneous 2C-like transition, with globally weakened chromatin loops. This included not only TAD boundary loops but also chromatin loops between ESC enhancers particularly super-enhancers and their neighboring gene promoters, thus leading to transcriptional down-regulation of many pluripotent genes controlled by ESC super-enhancers. Of note, down-regulation of pluripotent genes in 2CLCs were previously reported to take place only at the protein level but not the mRNA level (Eckersley-Maslin et al., 2016; Macfarlan et al., 2012). Nevertheless, a recent single-cell analysis of DUX-induced ESC to 2CLC transition showed a two-step reprogramming process with transcriptional inactivation of pluripotent genes as the first step (Fu et al., 2019), in agreement with our findings. The down-regulation of pluripotent genes at both mRNA and protein levels suggested a complex regulation of gene expression during ESC to 2CLC transition.

We further demonstrated a causal relationship between relaxed chromatin architecture and totipotency acquisition, by showing that disruption of chromatin loops in ESCs could facilitate 2C-like transition. We also observed formation of 2C enhancers during ESC to 2CLC transition. Importantly, up-regulated genes during the 2C-like transition tend to have ESC TAD boundaries separating them from neighboring 2C enhancers. Therefore, disruption of TAD boundary loops in ESCs may facilitate the contacts between 2C enhancers that are likely sporadically activated in ESCs and the promoters of nearby 2C genes, thus promoting their expression, which in turn fully activates the 2C-like transcriptional program. Consistently, a recent study showed that cohesin pre-depletion in donor cells could enhance SCNT efficiency at least partially through up-regulation of the minor ZGA genes in donor cells, many of which are also 2C-specific genes (Zhang et al., 2020).

In sum, we demonstrated that the spontaneous pluripotent to totipotent state transition in mouse ESCs not only coincide with global weakening of 3D genome conformation, but also can be promoted by disruption of chromatin loops. Further investigations are still required to elucidate the molecular mechanisms underlying the global weakening of higher-order chromatin structure during totipotency acquisition, as well as to fully understand how 2C-specific genes are activated upon disruption of chromatin loops.

## AUTHOR CONTRIBUTIONS

L.S. conceived and supervised the project; J.Y., J.G., and C.X. performed the experiments; Y.Z. and J.Y. analyzed the data; J.Y., Y.Z., and L.S. wrote the manuscript with inputs from L.Z. and J.C., and all authors discussed the results and commented on the manuscript.

## ACKNOWLEDGEMENTS

We thank Dr. Samuel Pfaff for the MERVL::tdTomato reporter, Dr. Benoit Bruneau for the CTCF-AID cell line, Dr. Yangming Wang and all members of the Shen laboratory for helpful discussions. This project was supported by National Natural Science Foundation of China (31871478, 32022023), National Key Research and Development Programs of China (2017YFC1001500), and Zhejiang Provincial Natural Science Foundation of China (LR18C060001).

## DATA AND SOFTWARE AVAILABILITY

The accession number for datasets reported in this study is GSE159623 (GEO). R scripts for data analysis are available upon request.

## EXPERIMENTAL PROCEDURES

### Cell lines and cell culture

The stable mouse ES cell line with the MERVL::tdTomato reporter (2C-E14) were generated as previously described (Macfarlan et al., 2012). Briefly, the MERVL::tdTomato reporter was transfected into E14 cells using Lipofectamine 2000 (11668030, Thermo Scientific), and single colonies were isolated. The 2C-CTCF-AID cell line was similarly generated by transfecting the MERVL::tdTomato reporter into the CTCF-AID cells. Mouse ES cells were cultured on 0.1% gelatin-coated plates with standard LIF/serum medium containing 15% FBS (VS500T, Ausbian), 1,000◻U/ ml mouse LIF (ESG1107, Millipore), 0.1 mM non-essential amino acids (11140, Gibco), 0.055◻mM β-mercaptoethanol (21985023, Gibco), 2 mM GlutaMAX (35050, Gibco), 1◻mM sodium pyruvate (11360, Gibco), and 100◻U/ml penicillin/streptomycin (15140, Gibco). For cell lines 2C-CTCF-AID cells, 500 μM indole-3-acetic acid (IAA, chemical analog of auxin, C3290, APExBIO) was used to induce CTCF degradation.

### sisHi-C

ESCs and 2CLCs were purified by FACS, and 100,000 cells were used for each Hi-C library. Cells were cross-linked for 10 min with 1% formaldehyde and quenched with 0.2 M glycine for 10 min at room temperature. The cross-linked cells were subsequently lysed in lysis buffer (10 mM Tris-HCl (pH 7.4), 10 mM NaCl, 0.5% NP-40, 0.1 mM EDTA and 1x proteinase inhibitor). The extracted nuclei were re-suspended with 10 μl 0.5% SDS and incubated at 62°C for 10 min. After adding 23 μl water and 5 μl 10% Triton X-100, and 15-min incubation at 37°C, genomic DNA was digested by adding 5 μl 10x NEB buffer 2 and 100 U of MboI. After overnight incubation at 37°C, the MboI enzyme was inactivated at 62°C for 20 min. Next, cohesive ends were filled in by adding 1.5 μl of 1 mM dCTP, 1.5 μl of 1 mM dGTP, 1.5 μl of 1 mM dTTP, 3.75 μl of 0.4 mM biotin-14-dATP and 2 μl (10 U) Klenow, and incubated at 37°C for 90 min. Subsequently, 39 μl water, 12 μl 10x T4 ligase reaction buffer, 7 μl 10% Triton X-100, 1.2 μl 100x BSA and 400 U T4 DNA ligase were added to start proximity ligation. The ligation reaction was placed at room temperature for 5.5h. After ligation, the crosslinking was reversed by 200 μg/ml proteinase K (AM2546, Thermo Fisher) and 12 μl 10% SDS at 55°C for 30 min, then add 13 μl 5 M NaCl, and incubate at 68°C overnight. DNA was purified and sheared to a length of ~400 bp. Point ligation junctions were pulled down by Dynabeads® MyOne™ Streptavidin C1 (65001, Thermo Fisher) according. The Hi-C library for Illumina sequencing was prepared after end preparation, adaptor ligation, PCR amplification and size selection. The final library was sequenced on the Illumina HiSeq X Ten platform with 150 paired-end mode.

### ULI-NChIP-seq

For ULI-NChIP-seq, 10,000 or 40,000 cells were used per reaction, and two or three replicates were performed. The ULI-NChIP procedure was performed as previously described (Brind’Amour et al., 2015). Two microgram of histone H3K27ac antibody (39133, Active Motif), histone H3K4me1 antibody (39297, Active Motif), histone H3K4me3 antibody (9727, Cell Signaling Technology), histone H3K27me3 antibody (9733, Cell Signaling Technology) or histone H3K9me3 antibody (39161, Active Motif) were used for each immunoprecipitation reaction. The sequence libraries were generated using the NEBNext Ultra II DNA Library Prep Kit for Illumina (E7645, New England Biolabs) following the manufacturer’s instructions. Barcoded libraries were pooled and sequenced on the Illumina HiSeq X Ten platform to generate 150 bp paired-end reads.

### ATAC-seq

The ATAC-seq libraries were prepared as previously described with minor modifications (Buenrostro et al., 2013). Briefly, samples were lysed in lysis buffer (10◻mM Tris-HCl (pH 7.4), 10◻mM NaCl, 3◻mM MgCl2 and 0.1% Igepal CA-630) for 10◻min on ice to prepare the nuclei. Immediately after lysis, nuclei were spun at 500 g for 5◻min to remove the supernatant. Nuclei were then incubated with the Tn5 transposome and tagmentation buffer at 37°C for 30◻min (TD501, Vazyme Biotech). After the tagmentation, the products were purified with 2.0x VAHTS DNA Clean Beads (N411, Vazyme Biotech). PCR was performed to amplify the library for 14 cycles using the following PCR conditions: 72°C for 3◻min; 98°C for 30◻s; and thermocycling at 98°C for 15◻s, 60°C for 30◻s and 72°C for 3◻min; following by 72°C 5◻min. After the PCR reaction, libraries were purified with the 1.2x VAHTS DNA Clean Beads. Barcoded libraries were pooled and sequenced on the Illumina HiSeq X Ten platform to generate 150 bp paired-end reads.

### ChIP-seq and ATAC-seq data analysis

Raw reads were trimmed to 50 bp and aligned to the mouse genome (mm9) using bowtie2 (version 2.3.4.1) with default parameters. Unmapped reads and PCR duplicates were removed. Peaks were called using MACS2 (version 2.1.1.20160309) with parameters “-q 0.05 --nomodel --nolambda --broad --extsize 300 -B --SPMR -g mm”. Average intensity profiles were generated using deepTools (version 2.5.4).

### RNA-seq and data analysis

RNA-seq libraries were prepared using the Smart-seq2 method as previously described (Picelli et al., 2013). Barcoded libraries were pooled and sequenced on the Illumina HiSeq X Ten platform in the paired-end mode. Raw reads were trimmed to 50 bp and mapped to the mouse genome (mm9) using TopHat (version 2.1.1) with default parameters. Only uniquely mapped reads were kept for downstream analysis. Gene counts were generated using HTSeq-count (version 0.9.1). For each sample, the gene count matrices were merged together and then the ‘Trimmed Mean of M values’ normalization (TMM) method (edgeR package in Bioconductor, version 3.5.0) was used to calculate the normalized expression.

### Hi-C data analysis

Raw reads were first processed with HiCPro (version 2.11.1). Briefly, raw reads were aligned to the mouse genome (mm9) using the Bowtie 2 end-to-end algorithm and ‘-very-sensitive’ option. To rescue the chimeric fragments spanning ligation junctions, the ligation site was detected and the 5’ fraction of the reads was aligned back to the reference genome. Unmapped reads, multiple mapped reads and singletons were then discarded. Pairs of aligned reads were then assigned to MboI restriction fragments. Read pairs from uncut DNA, self-circle ligation and PCR artefacts were filtered out, and the valid read pairs involving two different restriction fragments were used to build the contact matrix. Valid read pairs were then binned at a specific resolution by dividing the genome into bins of equal size. The binned interaction matrices were then normalized using the iterative correction method to correct for biases such as GC content, mappability and effective fragment length. For each chromosome, we obtained the expected Hi-C contact values by calculating the average contact intensity for all loci at a certain distance. We then transformed the normalized Hi-C matrix into an observed/expected (O/E) matrix by dividing each normalized observed value by its corresponding expected value. We identified A and B compartments and TADs from Hi-C data using cworld (Dekker lab, https://github.com/dekkerlab/cworld-dekker). To calculate PC1 values, we used the ‘‘matrix2compartment.pl’’ script with default parameters with contact maps binned at 500 kb resolution as input. Bins with positive or negative PC1 values were defined as A or B compartment, respectively. The compartment strength was then calculated as (AA+BB)/2AB as previously described (Flyamer et al., 2017). To calculate insulation scores, we used the “matrix2insulation.pl” script with default parameters at 10 kb resolution. Insulation scores were further normalized by dividing each bin’s insulation score by the average scores in the nearest 2 Mb region, log2-transformed, and multiplied by −1. The “insulation2tads.pl” script was used with default parameters to identify stable TADs as previously described (Ke et al., 2017). Directionality index (DI) scores were calculated using a previously described pipeline (Dixon et al., 2012). The relative frequency of inter-TAD interactions was calculated from the number of interactions between 2 TADs divided by the total number of cis-interactions in the corresponding chromosome. Aggregate Hi-C contact maps were generated by extracting subsets from the OE matrix (these can be either single regions along the diagonal, or region pairs corresponding the matrix segments off the diagonal) and averaging all resulting sub-matrices. If the sub-matrices were of different size, they were interpolated to a fixed size. TAD strength was calculated as the mean of values in the OE matrix in the TAD region. Loop strength was calculated as the mean of values in the OE matrix in the nearest 10 regions of both anchors. Global contact decay curves were plotted as previously described (Olivares-Chauvet et al., 2016). In order to better visualize differences between conditions, we calculated the distribution of the Hi-C contacts as a simple contact probability (sum of the observed counts per log2 bin, divided by total observed contacts, without normalizing for the bin size).

### siRNA transfection

ESCs were transfected with siRNAs (Gene Pharma) using the DharmaFECT 1 (T-2001, Dharmacon) reagent following the manufacturer’s instructions. The control siRNA (siNC) was derived from sequences in the *Caenorhabditis elegans* genome and does not target any mammalian genes. Cells were collected for RNA extraction 2 days after transfection.

### RNA isolation, and RT-qPCR

Total RNA was isolated using RNeasy Plus Mini Kit (74134, Qiagen), cDNA was generated using HiScript II Q RT Super Mix with gDNA wiper (R223, Vazyme Biotech), and qPCR was performed using the AceQ qPCR SYBR Green Master Mix (Q111, Vazyme Biotech). Relative quantification was performed using the comparative Ct method with normalization to *Actin*. Primer sequences are available in Table S2.

### Statistical analysis

Statistical analyses were performed with the R software. Statistics were calculated using the two-tailed Wilcox signed rank test (for paired samples) unless indicated otherwise.

## SUPPLEMENTAL INFORMATION

**Document S1. Figures S1-S4.**

**Table S1. Sequencing information summary, related to Figure 1–3.** Sequencing summary for Hi-C libraries, RNA-seq libraries, ATAC-seq libraries and ChIP-seq libraries.

**Table S2. Sequences for real-time qPCR primers and siRNAs, related to Figure 2 snd 4.** Sequence information of forward and reverse primers used for pluripotent genes and various ZGA transcripts in qPCR. Sequence information of sense and antisense siRNA strands for knocking down *Ctcf*, *Smc1a*, *Smc3*, *Rad21*, or *Yy1*.

