## Supplementary material for "Relaxed 3D genome conformation facilitates the pluripotent to totipotent state transition in embryonic stem cells": Document S1

Figure S1

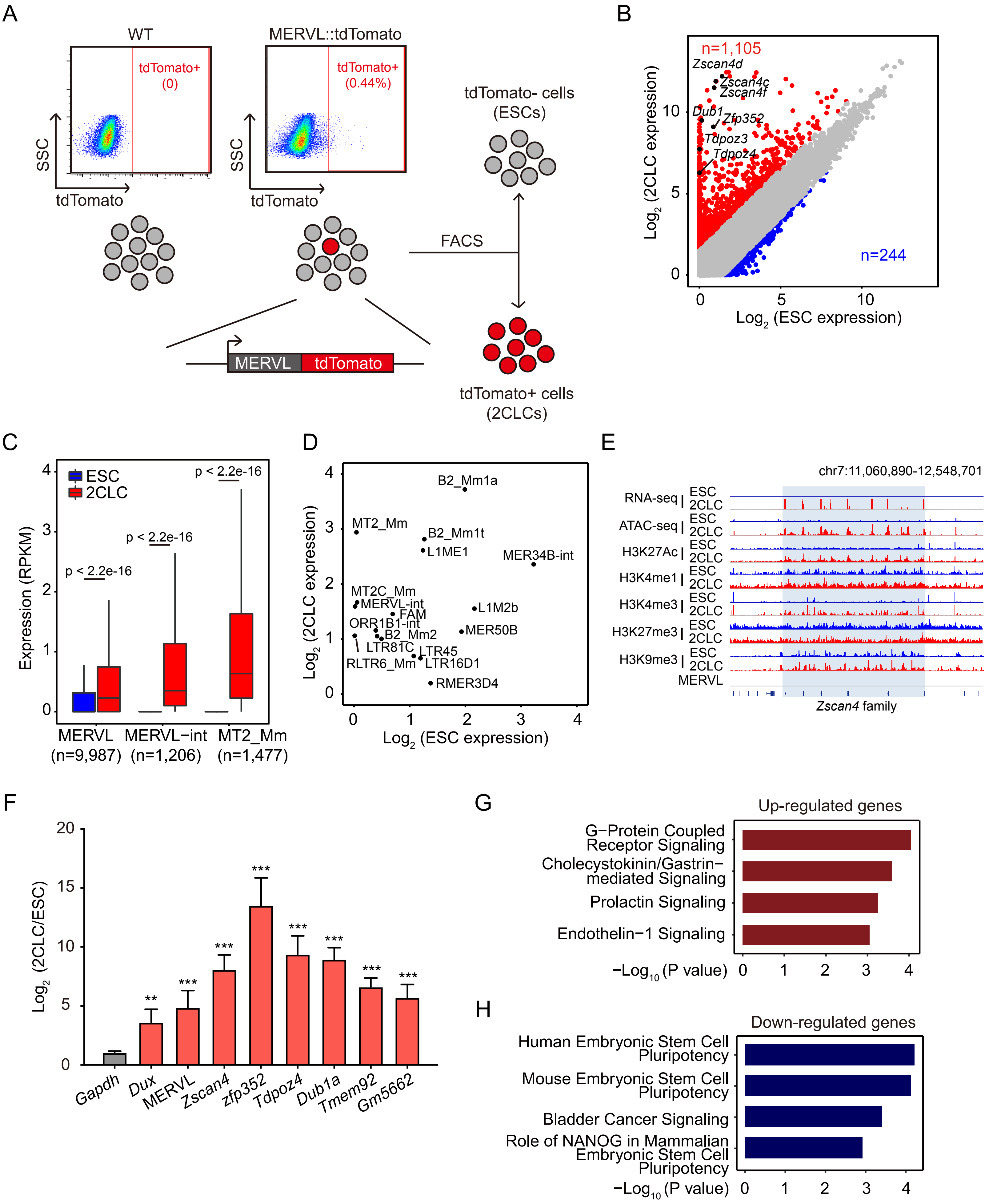

A
WT
MERVL::tdTomato
tdTomato+
(0)
tdTomato+
(0.44%)
SSC
tdTomato
tdTomato- cells
(ESCs)
FACS
MERVL
tdTomato
tdTomato+ cells
(2CLCs)
B
n=1,105
Zscan4d
Zscan4c
Zscan4f
Dub1
Zfp352
Tdpoz3
Tdpoz4
n=244
Log2 (2CLC expression)
Log2 (ESC expression)
C
ESC
2CLC
p < 2.2e-16
Expression (RPKM)
MERVL
(n=9,987)
MERVL−int
(n=1,206)
MT2_Mm
(n=1,477)
D
B2_Mm1a
MT2_Mm
B2_Mm1t
L1ME1
MER34B-int
MT2C_Mm
MERVL-int
FAM
ORR1B1-int
B2_Mm2
LTR81C
RLTR6_Mm
LTR45
LTR16D1
L1M2b
MER50B
RMER3D4
Log2 (2CLC expression)
Log2 (ESC expression)
E
chr7:11,060,890-12,548,701
RNA-seq
ATAC-seq
H3K27Ac
H3K4me1
H3K4me3
H3K27me3
H3K9me3
ESC
2CLC
MERVL
Zscan4 family
F
Log2 (2CLC/ESC)
Gapdh
Dux
MERVL
Zscan4
zfp352
Tdpoz4
Dub1a
Tmem92
Gm5662
G
Up-regulated genes
G−Protein Coupled Receptor Signaling
Cholecystokinin/Gastrin−mediated Signaling
Prolactin Signaling
Endothelin−1 Signaling
−Log10 (P value)
H
Down-regulated genes
Human Embryonic Stem Cell Pluripotency
Mouse Embryonic Stem Cell Pluripotency
Bladder Cancer Signaling
Role of NANOG in Mammalian Embryonic Stem Cell Pluripotency
−Log10 (P value)

**Figure S1. RNA-seq validates the fidelity of the purified 2CLCs**

(A) Schematic representation of 2CLC purification.

(B) Scatter plot comparing the gene expression between ESC and 2CLC. The criteria for gene expression changes are log2(FC) > 1.5.

(C) Box plot showing expression of MERVL, MERVL-int, and MT2_Mm in ESC and 2CLC.

(D) Scatter plot of transcriptional level of repeats in ESC and 2CLC.

(E) Genome browser view showing RNA-seq, ATAC-seq, H3K27Ac, H3K4me1, H3K4me3, H3K27me3, and H3K9me3 signals at *Zscan4* and MERVL locus in ESC and 2CLC. Blue box = *Zscan4* locus.

(F) Relative expression levels of 2cell-specific elements in 2CLCs. Data are normalized to *Actin* and are presented as mean ± SD, $^{**}p < 0.01$, $^{***}p < 0.001$ (Multiple *t* tests). *Gapdh* serves as a control.

(G) Ingenuity pathway analysis of up-regulated genes in 2CLCs.

(H) Ingenuity pathway analysis of down-regulated genes in 2CLCs.

### Figure S2

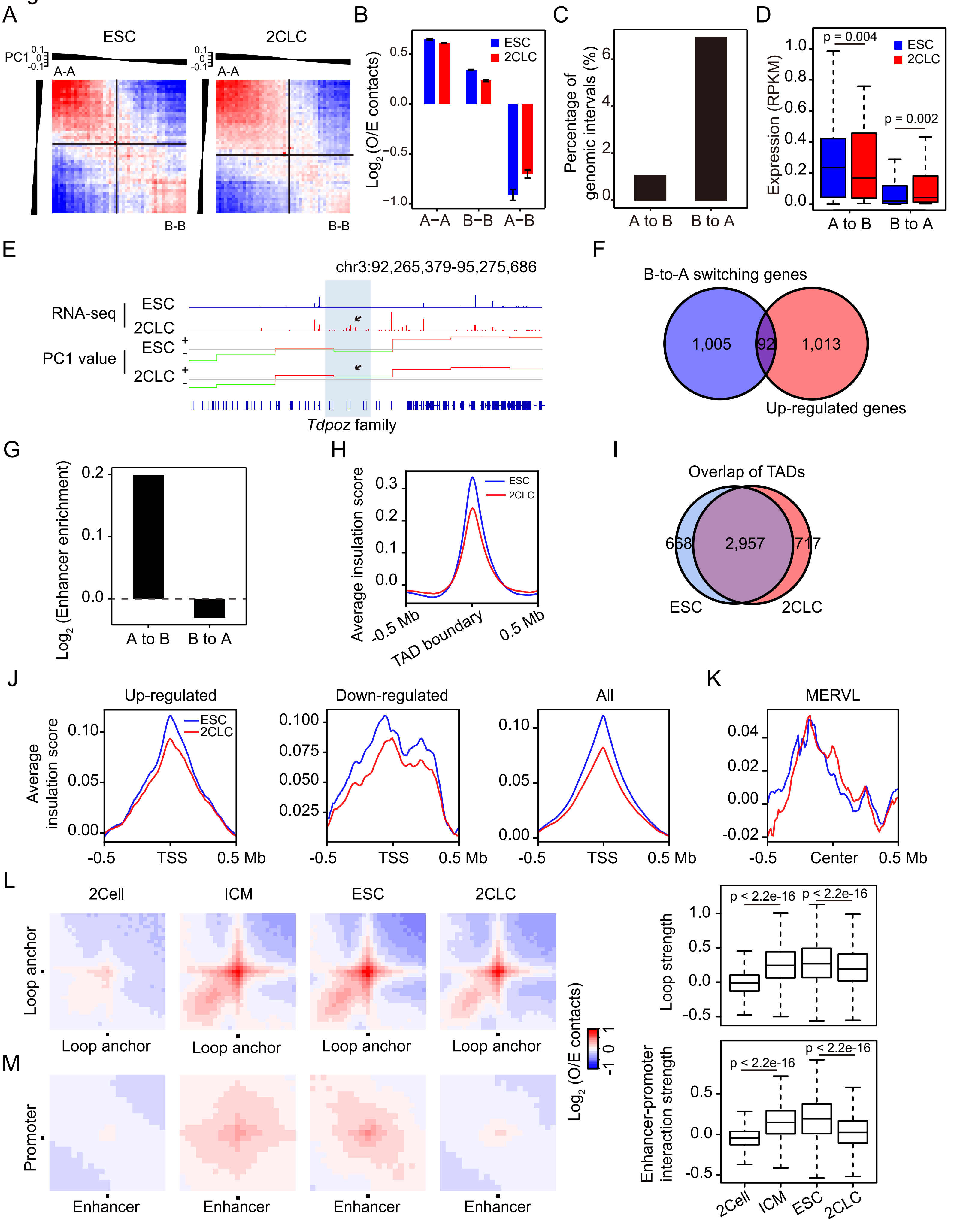

A
ESC
2CLC
PC1
A-A
B-B
B
Log2 (O/E contacts)
ESC
2CLC
A−A
B−B
A−B
C
Percentage of genomic intervals (%)
A to B
B to A
D
Expression (RPKM)
p = 0.004
p = 0.002
A to B
B to A
E
chr3:92,265,379-95,275,686
RNA-seq
PC1 value
Tdpoz family
F
B-to-A switching genes
1,005
92
1,013
Up-regulated genes
G
Log2 (Enhancer enrichment)
A to B
B to A
H
Average insulation score
-0.5 Mb
TAD boundary
0.5 Mb
I
Overlap of TADs
668
2,957
717
ESC
2CLC
J
Up-regulated
Down-regulated
All
Average insulation score
-0.5
TSS
0.5 Mb
K
MERVL
Center
L
2Cell
ICM
ESC
2CLC
Loop anchor
Loop strength
p < 2.2e-16
Log2 (O/E contacts)
M
Promoter
Enhancer
Enhancer-promoter interaction strength

**Figure S2. 3D chromatin structures in ESCs, 2CLCs and early embryos.**

(A) Hi-C contact maps between pairs of 500-kb loci arranged by their PC1 values (shown on top and left) across chromosome 1.

(B) Bar plot showing O/E contact strength between domains from the same ("A" versus "A" or "B" versus "B") and different ("A" versus "B") compartment in ESC and 2CLC. Values represent means of two biological replicates with ends of error bars corresponding to individual data points.

(C) Bar plot showing the percentage of genomic intervals which display A/B compartment switch during ESC to 2CLC transition.

(D) Box plot showing the transcriptional level of regions display A/B compartment switch.

(E) Genome browser view showing RNA-seq signal and PC1 values at *Tdpoz* locus in ESC and 2CLC. Green = B compartment; red = A compartment; Blue box = *Tdpoz* locus.

(F) Venn diagram showing the overlap of B-to-A switching genes and up-regulated genes in 2CLC.

(G) Bar plot showing log2 ratio of ESC enhancers to control regions located in A/B compartment switches regions. Equal number of random regions were used as control.

(H) Average insulation score in a 0.5 Mb region centered on TAD boundaries.

(I) Venn diagram showing the overlap of TADs identified in ESC and 2CLC.

(J-K) Average insulation score around TSS of up-regulated, down regulated and all genes (J), and MERVL (K) in ESC and 2CLC (±0.5 Mb).

(L) Aggregate Hi-C contact maps between pairs of loop anchors in 2-cell embryo (2Cell), ICM embryo (ICM), ESC, and 2CLC (left). Box plot showing loop strength in 2Cell, ICM, ESC, and 2CLC (right).

(M) Aggregate Hi-C contact maps between ESC enhancer-promoter pairs in 2-cell embryo (2Cell), ICM embryo (ICM), ESC, and 2CLC (left). Box plot showing loop strength in 2Cell, ICM, ESC, and 2CLC (right).

Figure S3

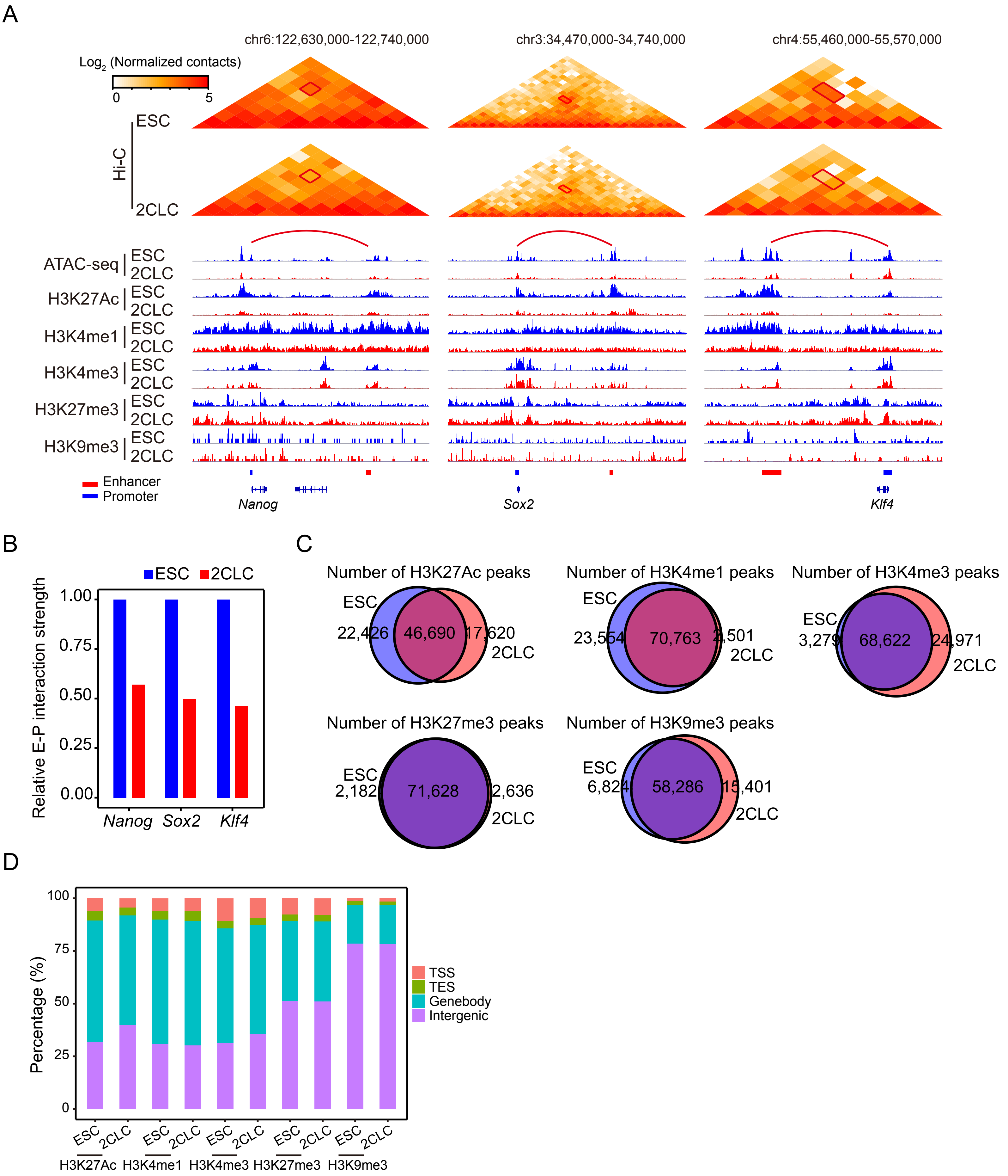

A
chr6:122,630,000-122,740,000
chr3:34,470,000-34,740,000
chr4:55,460,000-55,570,000
Log2 (Normalized contacts)
0
5
Hi-C
ESC
2CLC
ATAC-seq
H3K27Ac
H3K4me1
H3K4me3
H3K27me3
H3K9me3
ESC
2CLC
Enhancer
Promoter
Nanog
Sox2
Klf4
B
ESC
2CLC
Relative E-P interaction strength
1.00
0.75
0.50
0.25
0.00
Nanog
Sox2
Klf4
C
Number of H3K27Ac peaks
ESC
22,426
46,690
17,620
2CLC
Number of H3K4me1 peaks
ESC
23,554
70,763
2,501
2CLC
Number of H3K4me3 peaks
ESC
3,279
68,622
24,971
2CLC
Number of H3K27me3 peaks
ESC
2,182
71,628
2,636
2CLC
Number of H3K9me3 peaks
ESC
6,824
58,286
15,401
2CLC
D
Percentage (%)
100
75
50
25
0
TSS
TES
Genebody
Intergenic
ESC
2CLC
H3K27Ac
H3K4me1
H3K4me3
H3K27me3
H3K9me3

**Figure S3. Loss of enhancer-promoter interactions of pluripotent genes in 2CLCs**

(A) Hi-C contact maps near *Nanog*, *Sox2*, and *Klf4* (top). Genome browser tracks of ATAC, H3K27Ac, H3K4me1, H3K4me3, H3K27me3, and H3K9me3 ChIP-seq signals in the corresponding region (bottom). An arc representing an enhancer-promoter interaction. The signal in the outlined pixels was used to quantify the change in enhancer-promoter interaction strength between ESC and 2CLC.

(B) Bar plot showing the relative enhancer-promoter interaction strength in ESC and 2CLC at the *Nanog*, *Sox*, and *Klf4* locus, which are outlined at (C). E-P interaction = enhancer-promoter interaction.

(C) Venn diagram showing the overlap of H3K27Ac, H3K4me1, H3K4me3, H3K27me3, and H3K9me3 ChIP-seq peaks identified in ESC and 2CLC.

(D) Bar plot showing the genomic distribution of H3K27Ac, H3K4me1, H3K4me3, H3K27me3, and H3K9me3 ChIP-seq peaks in ESC and 2CLC.

Figure S4

A

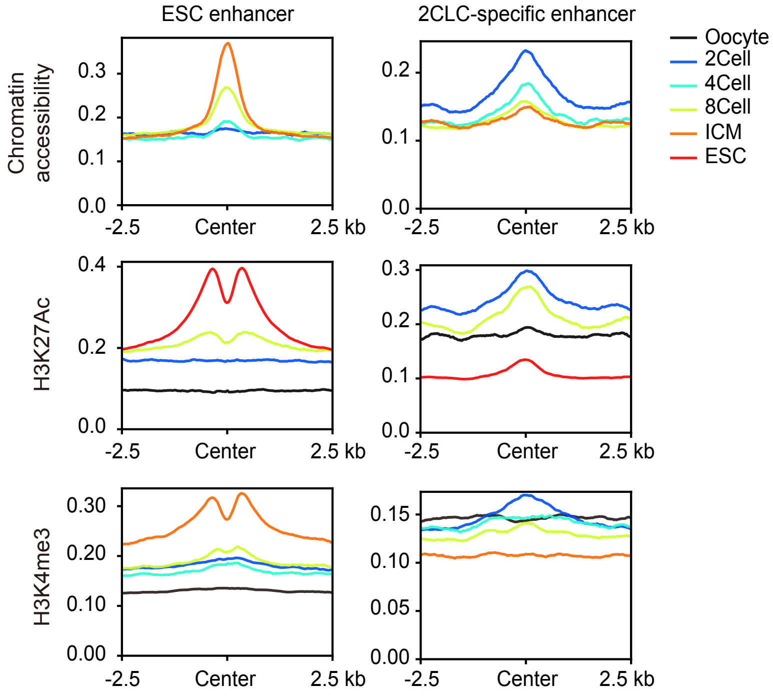
ESC enhancer
2CLC-specific enhancer
Oocyte
2Cell
4Cell
8Cell
ICM
ESC
Chromatin accessibility
H3K27Ac
H3K4me3
0.0
0.1
0.2
0.3
0.4
0.00
0.05
0.10
0.15
0.20
0.30
-2.5
Center
2.5 kb

B

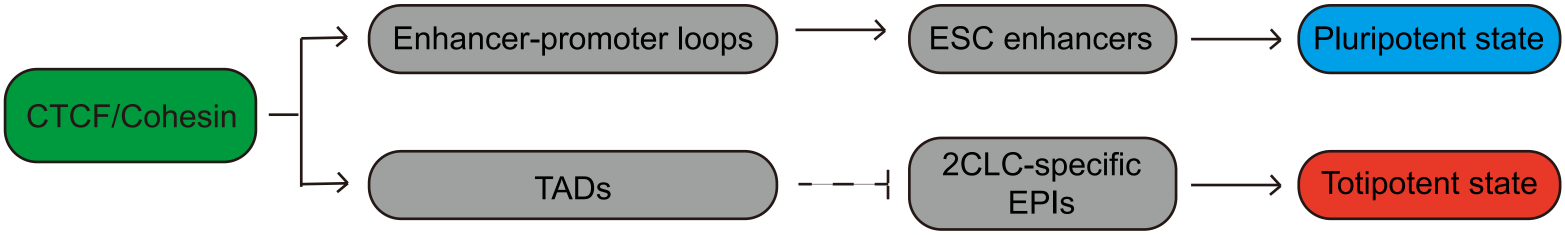
CTCF/Cohesin
Enhancer-promoter loops
TADs
ESC enhancers
2CLC-specific EPIs
Pluripotent state
Totipotent state

**Figure S4. Shifts between ESC enhancers and 2C enhancers in 2-cell embryos**

(A) Average ATAC, H3K27Ac, and H3K4me3 signals in 2.5 kb region centered on ESC enhancers and 2C enhancers in oocyte, 2-cell, 4-cell, 8-cell, ICM embryos, and ESCs.

(B) A model showing that disruption of enhancer-promoter loops and TADs facilitates the ESC to 2CLC transition.
